# Utilizing Evolutionary Conservation to Detect Deleterious Mutations and Improve Genomic Prediction in Cassava

**DOI:** 10.1101/2022.09.16.508169

**Authors:** Evan M Long, M. Cinta Romay, Edward S. Buckler, Kelly R Robbins

## Abstract

Cassava (*Manihot esculenta*) is an annual root crop which provides the major source of calories for over half a billion people around the world. Since its domestication ∼10,000 years ago, cassava has been largely clonally propagated through stem cuttings. Minimal sexual recombination has led to an accumulation of deleterious mutations made evident by heavy inbreeding depression. To locate and characterize these deleterious mutations, and to measure selection pressure across the cassava genome, we aligned 52 related Euphorbiaceae and other related species representing millions of years of evolution. With single base-pair resolution of genetic conservation, we used protein structure models, amino acid impact, and evolutionary conservation across the Euphorbiaceae to estimate evolutionary constraint. With known deleterious mutations, we aimed to improve genomic evaluations of plant performance through genomic prediction. We first tested this hypothesis through simulation utilizing multi-kernel GBLUP to predict simulated phenotypes across separate populations of cassava. Simulations showed a sizable increase of prediction accuracy when incorporating functional variants in the model when the trait was determined by <100 quantitative trait loci (QTL). Utilizing deleterious mutations and functional weights informed through evolutionary conservation, we saw improvements in genomic prediction accuracy that were dependent on trait and prediction. We showed the potential for using evolutionary information to track functional variation across the genome, in order to improve whole genome trait prediction. We anticipate that continued work to improve genotype accuracy and deleterious mutation assessment will lead to improved genomic assessments of cassava clones.

## Introduction

### Cassava

Cassava (Manihot esculenta) is a root crop that is clonally propagated and grown widely in the tropical regions of Africa, Asia, and South America. It is estimated that cassava is a major caloric source for almost half a billion people around the world(Parmar et al., 2017; Ferguson et al., 2019). Although it is naturally an outcrossing perennial, it has been clonally propagated and grown as an annual since its domestication between 5,000-10,000 years ago (Wang et al., 2014). During the colonial era it was also brought to Africa, where today it is valued for its ability to grow with minimal inputs in marginally fertile lands.

### Deleterious Mutations

Many generations of clonal propagation have caused cassava to accumulate genetic load that inhibits its potential crop performance. This genetic load is most apparent in the heavy inbreeding depression exhibited in cassava, as observed through low performance of selfed offspring (Rojas et al., 2009; de Freitas et al., 2016). Studies have shown that this genetic load is present as deleterious recessive mutations that are masked by heterozygosity which can be maintained through the clonal propagation (Ramu et al., 2017). With minimal sexual reproduction these deleterious mutations are maintained (McKey et al., 2010) and inhibit current breeding efforts to improve cassava performance (de Freitas et al., 2016).

### Genetic Load and Selection

Plant breeders have worked on various methods to detect and manage genetic load throughout history. Many crop species exist as polyploids, which enables them to more easily mask recessive deleterious mutations responsible for genetic load (van de Peer et al., 2021). Hybrid crop breeding has been another common method of applying strong selection pressures by selecting on inbred lines (Labroo et al., 2021), eliminating the possibility of recessive deleterious mutations. Some crops with similar high inbreeding depression to cassava, like potato, have made recent efforts to breed with inbred diploids (Bachem et al., 2019), however the deleterious mutations targeted by this methodology reduce plant viability.

During the past decade, plant breeders have seen the emergence of methodical application of genotyping and genomic selection as a method to improve breeding selections and leverage understanding of genomic information. Genomic selection, which uses genome markers and a phenotyped training population to predict unobserved offspring performance, can decrease selection cycle time and improve selection accuracy. Efforts have been made to improve genomic selection by using causative knowledge, however understanding the true causative elements in the genome is not a trivial exercise. Many studies have shown that including genome-wide association (GWA) hits in prediction can breakdown when predicting unrelated material (Cheruiyot et al., 2022), indicating population specific quantitative trait locus (QTL) or a misinterpretation of a variant as causative, when it is only in high linkage disequilibrium (LD) with the causative variant (Cheruiyot et al., 2022). For cassava, an ideal genomic annotation would explain underlying causative elements, while being consistent across populations structures.

Regarding genetic load, evolutionary conservation has shown to be an effective method to assess deleterious mutations and explain functional variation (Xiang et al., 2019) in a population agnostic manner. Multiple studies in crops such as maize (Yang et al., 2016; Ramstein and Buckler, 2022), sorghum (Valluru et al., 2019; Lozano et al., 2021), and barley (Kono et al., 2019) have demonstrated potential benefits for detecting and using deleterious mutations in genomic prediction. The potential benefit of understanding these deleterious mutations in cassava will be limited by the absolute number of mutations and how much variation of agronomic traits they each explain.

### Evaluating Deleterious Mutations in Genomic Prediction

The purpose of this study is first, to identify likely deleterious mutations in cassava, and second to evaluate their potential impact on genomic prediction for the goal of improving future breeding selections. We sequenced, assembled, and gathered 52 genomes from species that all shared ancestry within the last 50 million years in order to score conservation and detect deleterious mutations.

We designed an experiment that uses evolutionary information to augment genomic predictions within and across two different populations of 1048 cassava clones present in two different breeding programs in Sub-Saharan Africa, the International Institute of Tropical Agriculture (IITA), Ibadan, Nigeria, and the National Crops Resources Research Institute (NaCRRI), Namulonge, Uganda. By performing phenotype simulations using real genotypic data and generating genomic predictions with known, simulated QTL, we first evaluated the best possible benefit of including causative information in our genomic predictions under different scenarios. We then used genomic and phenotypic data from these cassava clones to test genomic predictions, while including various functional annotations based on deleterious mutations.

## Results

### Phenotype Simulation

To validate our methodology and guide our expectations we performed genomic predictions using simulated phenotypes on 1048 cassava clones originating from IITA and NaCRRI breeding programs. These simulations represent some best-case scenarios for genomic prediction, where all QTL and their effect sizes are known.

The simulated QTL effects represent a suite of different genetic architectures ranging from highly complex genetic traits controlled by thousands of small effect QTL to oligogenic traits controlled by a handful of large effect QTL. These genetic architectures are represented by the proportion of the 66k variants simulated as causative QTL (Fig. 1). We modeled dominance at each QTL in order to more closely match our empirical scenario in cassava, where genetic load due to recessive deleterious alleles are expected to affect many agronomic, fitness related, traits (Bosse et al., 2019).

**Figure 1.**
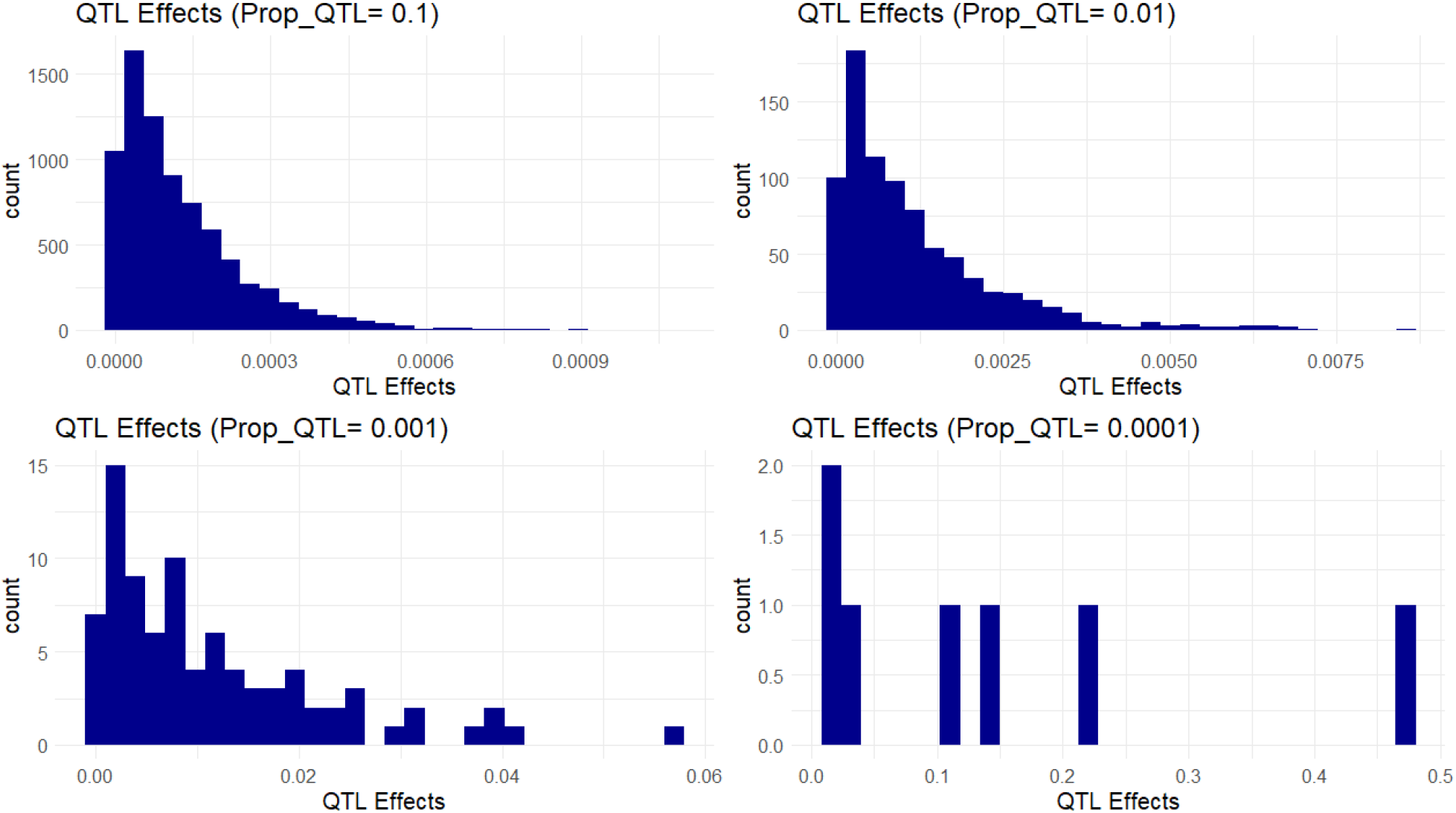
Simulated QTL Effects. Histograms show count of QTL effects in one example simulation. Each facet shows a genetic architecture with different proportions of the markers acting as QTL (resulting in ∼ 6600, 660, 66, and 6 QTL on average). The x-axis represents the positive effect of carrying the ancestral allele at a given QTL.

### Genomic Prediction with Simulated Phenotypes

Once QTL effects were modeled, we then calculated phenotypes for each of the 1048 clones (Sup Fig 1), where a positive effect is attributed to the ancestral allele. To Evaluate the effect of QTL structure, prediction model, and population, we performed genomic predictions. For all predictions shown, our cross-population and within-population predictions follow the representative schema (Fig 2). Cross-population prediction accuracy is calculated by masking all phenotypes in one population and predicting using the other, then calculating the correlation between the true phenotype and the predicted phenotype. Within-population prediction accuracy is calculated similarly, using a 10-Fold prediction scheme where phenotypes in 10% of a population are masked and predicted by the other 90%.

**Figure 2.**
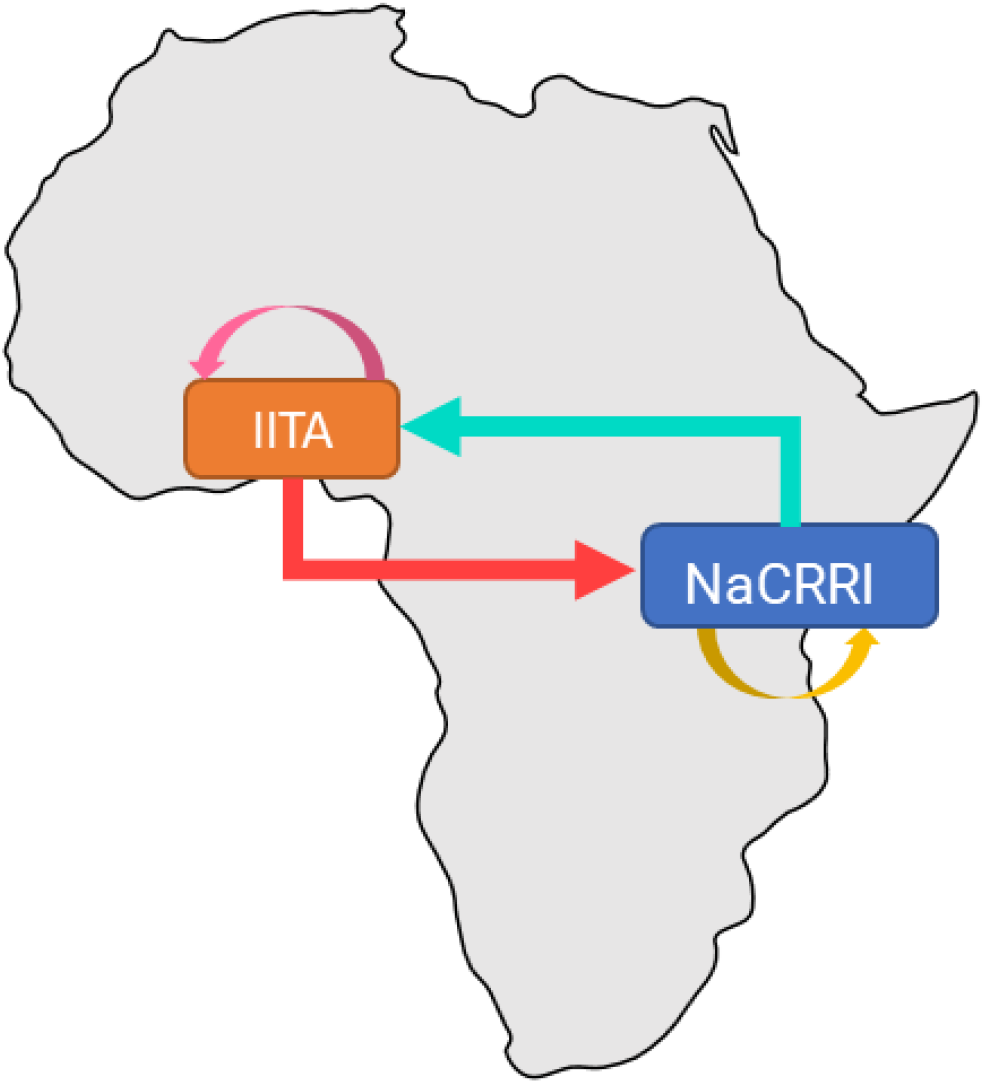
Genomic Prediction Schemes. Arrows represent genomic prediction schemes, where the base of each arrow represents the clones used to train the model and the tip represents the clones whose phenotypes were masked and then predicted. Arrows correspond to genomic predictions as follow “IITA_CV” (pink), “NaCRRI_CV” (yellow), “IITA->NaCRRI” (red), and “NaCRRI->IITA” (teal)

We saw a marked increase in prediction accuracy when including the QTL information into the prediction model only when the trait was controlled by less than around 100 QTL (Fig 3 bottom panels). Complex traits that are controlled by many small effect QTL across the genome show no increase in prediction accuracy with the inclusion of causative information (Fig 3 top panels). For traits with an intermediate number of QTL (Fig 3 bottom-left panel), the improvements in prediction accuracy are further increased by weighting the QTL information by their relative effect sizes, while under narrow genetic architectures most of the functional variation is captured without weighting. While the improvements are visible in both cross-population and within-population predictions, the improvements show some evidence of being more pronounced in cross-populations scenarios.

**Figure 3.**
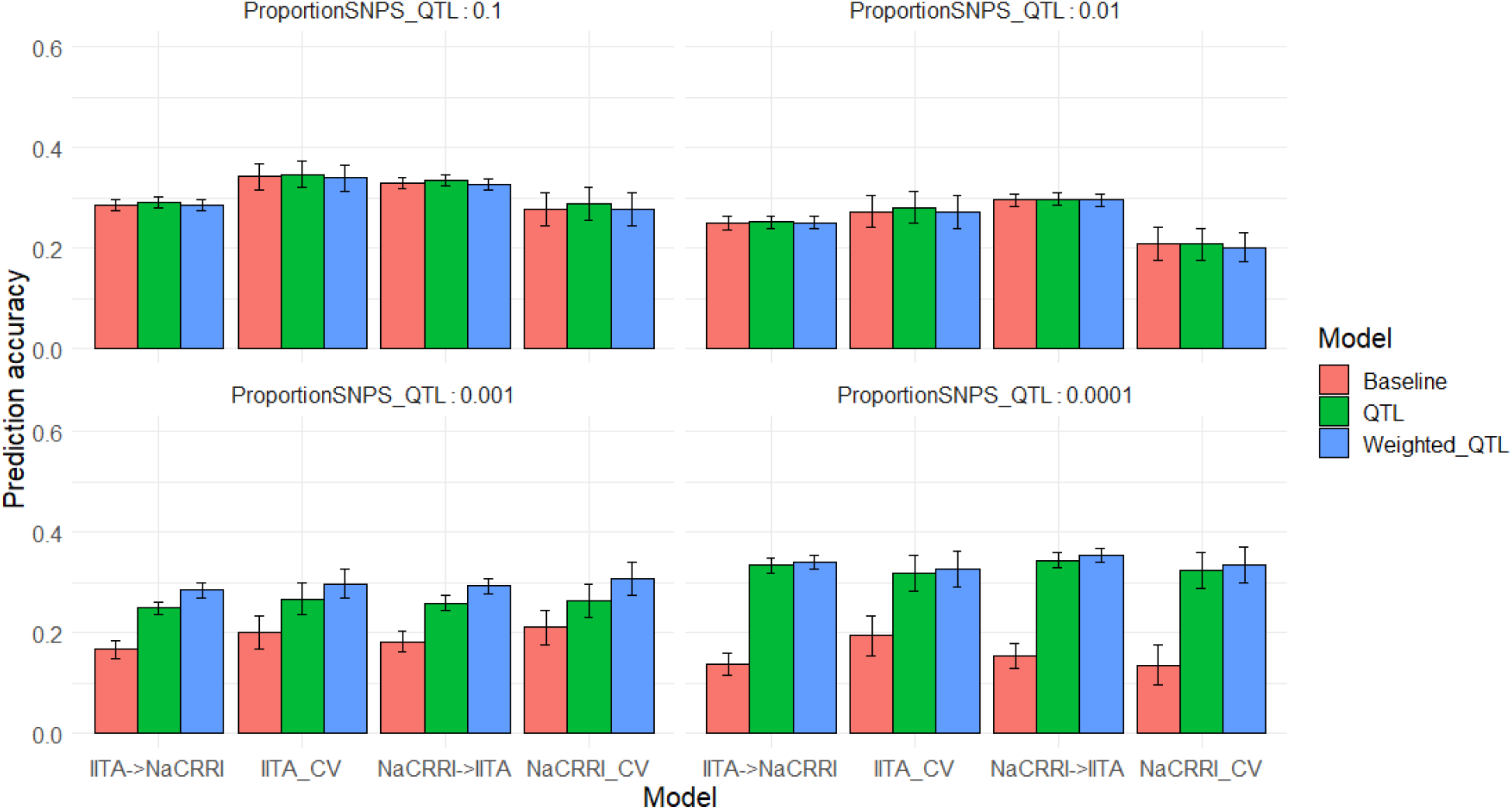
Genomic Prediction Accuracies with Simulated QTL. Prediction accuracies are shown on the y-axis as the correlation between predicted and true breeding values. The x-axis delineates the prediction scenario being tested. Barplot color corresponds to the genomic information used in the prediction model. Error bars represent a 95% confidence interval for simulations.

## Deleterious Mutations

We used evolutionary conservation and predicted protein mutation effects to classify the deleterious effects of 66k nonsynonymous SNPs segregating in the two target populations.

Firstly, we used the intersection of baseml evolutionary rate and SIFT deleterious scores to classify 2,210 deleterious sites that are segregating in both cassava populations (Fig. 4). Deleterious burden for each clone was then calculated as the number of derived alleles at these sites. We separated this deleterious burden into homozygous and heterozygous genetic load. Genome wide association for all nonsynonymous sites as well as the deleterious sites was performed on fresh root yield and dry matter percentage traits, and some loci passed Bonferroni significance testing for fresh root yield (Sup Fig 4&5).

**Figure 4.**
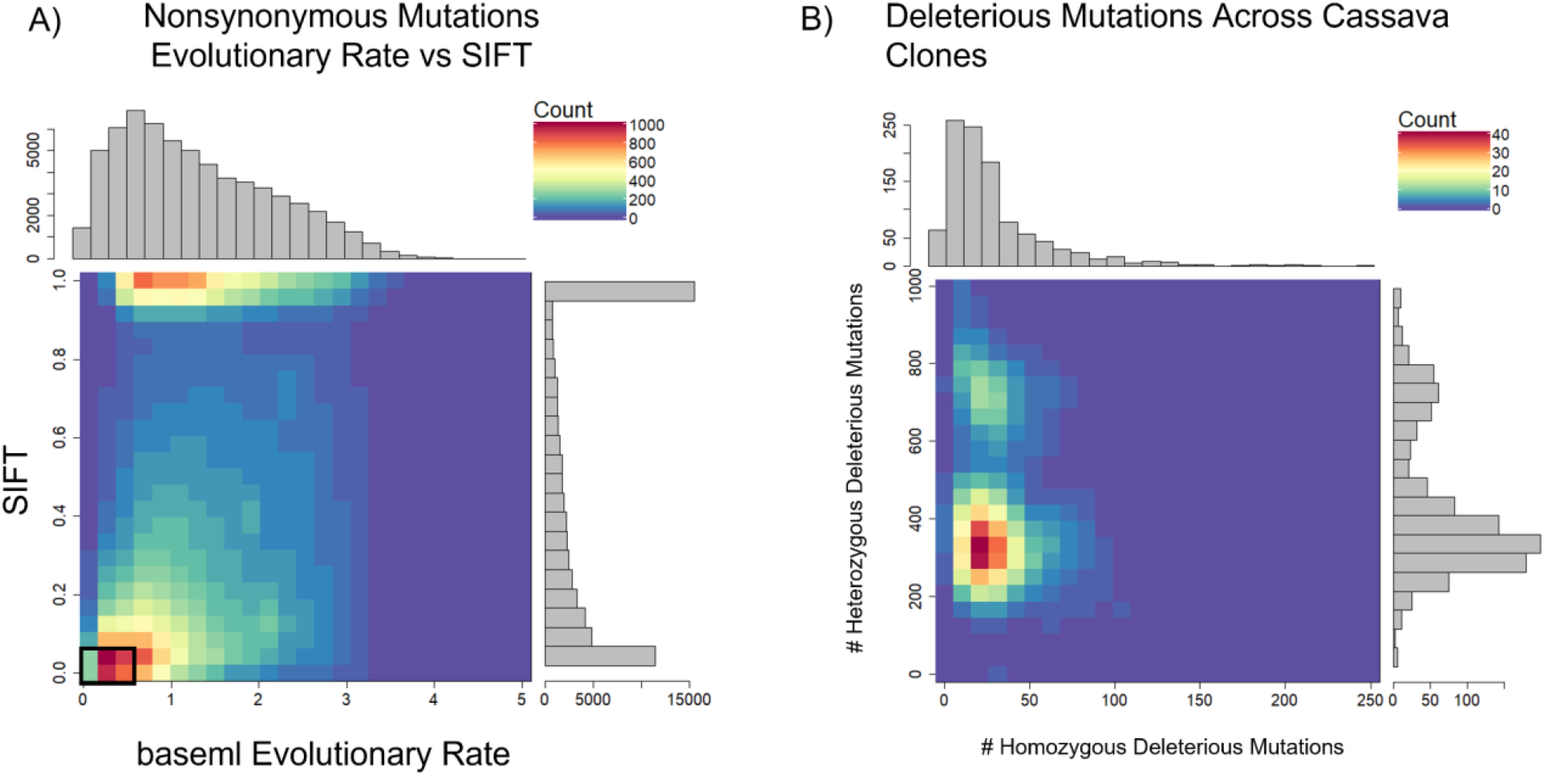
Defining Deleterious Mutations. A) baseml evolutionary rate is plotted against SIFT scores. Deleterious mutations were classified as derived alleles at those sites with a baseml evolutionary rate < 0.5 and a SIFT score < 0.05 (Black box). B) Distribution of homozygous and heterozygous deleterious mutations across 1048 cassava clones

**Figure 5.**
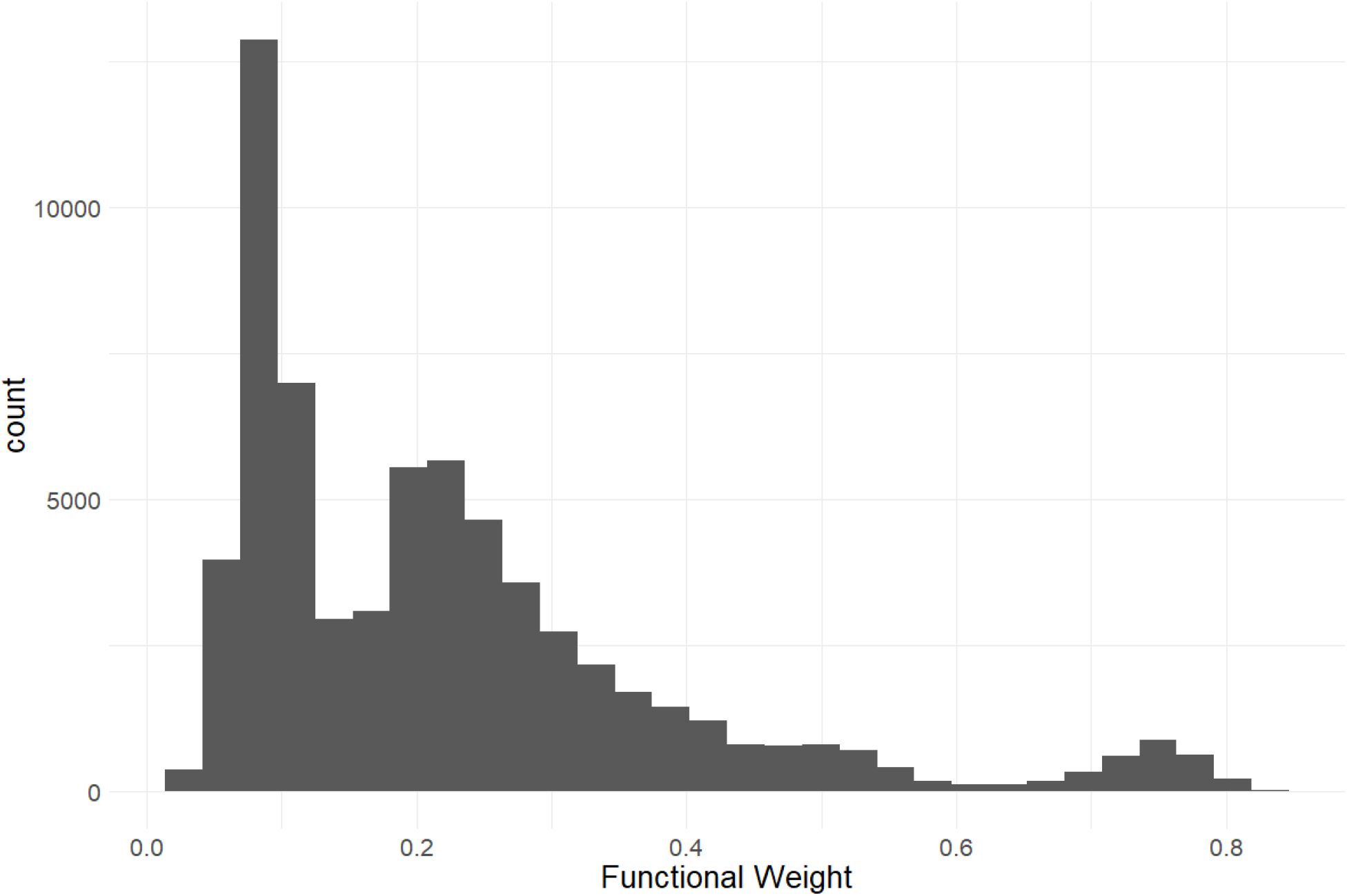
Predicted Functional Weights. Histogram of functional weights produced through RandomForest prediction of conservation for nonsynonymous variant sites. High functional weights correspond to highly conserved sites where nonsynonymous mutations are predicted to have large functional effects.

Secondly, we leveraged a RandomForest prediction model to weight the functional importance of the nonsynonymous mutations. The RandomForest model which used baseml evolutionary rate as a response, while leveraging protein annotation scores SIFT (Ng and Henikoff, 2003) and UniRep (Alley et al., 2019) as predictors. This prediction produces a score between 0-1, a quantitative weight for the functional importance of each amino acid residue altered by mutations at the nonsynonymous sites (Fig 5).

### Genomic Prediction Utilizing Functional Annotation

With deleterious mutations and functional weights for the segregating nonsynonymous sites, we mirrored the genomic predictions that we previously performed using simulated phenotypes, only this time using real data collected on the 1048 cassava clones.

We predicted two different traits common in cassava breeding trials, fresh root yield and dry matter percentage, using the same cross-population and within-population scenarios previously shown. Multiple genomic prediction models were tested to evaluate the value of including the functional annotations.

Our two examples of a baseline prediction, where no functional information is present, are genomic prediction using the input marker data set and a genome-wide imputed dataset. In predicting fresh root yield, our results show that imputation alone does not improve cross-population prediction accuracy, however it does show some positive effect on within-population prediction (Fig. 6). However, when including only imputed, segregating, non-synonymous variants, the prediction accuracy in cross-population predictions does increase over the two baseline models. Finally, we observed a further increase in prediction accuracy when weighting the non-synonymous variants and including derived genetic load from the deleterious mutations for both the cross-population predictions of fresh root yield and for within-population predictions in among the NaCRRI clones (Fig 6, Sup Fig 6). For genomic prediction of cassava tuber dry matter percentage, we observed mostly negative or neutral effects of imputation and inclusion of deleterious annotations (Fig 7, Sup Fig. 7).

**Figure 6.**
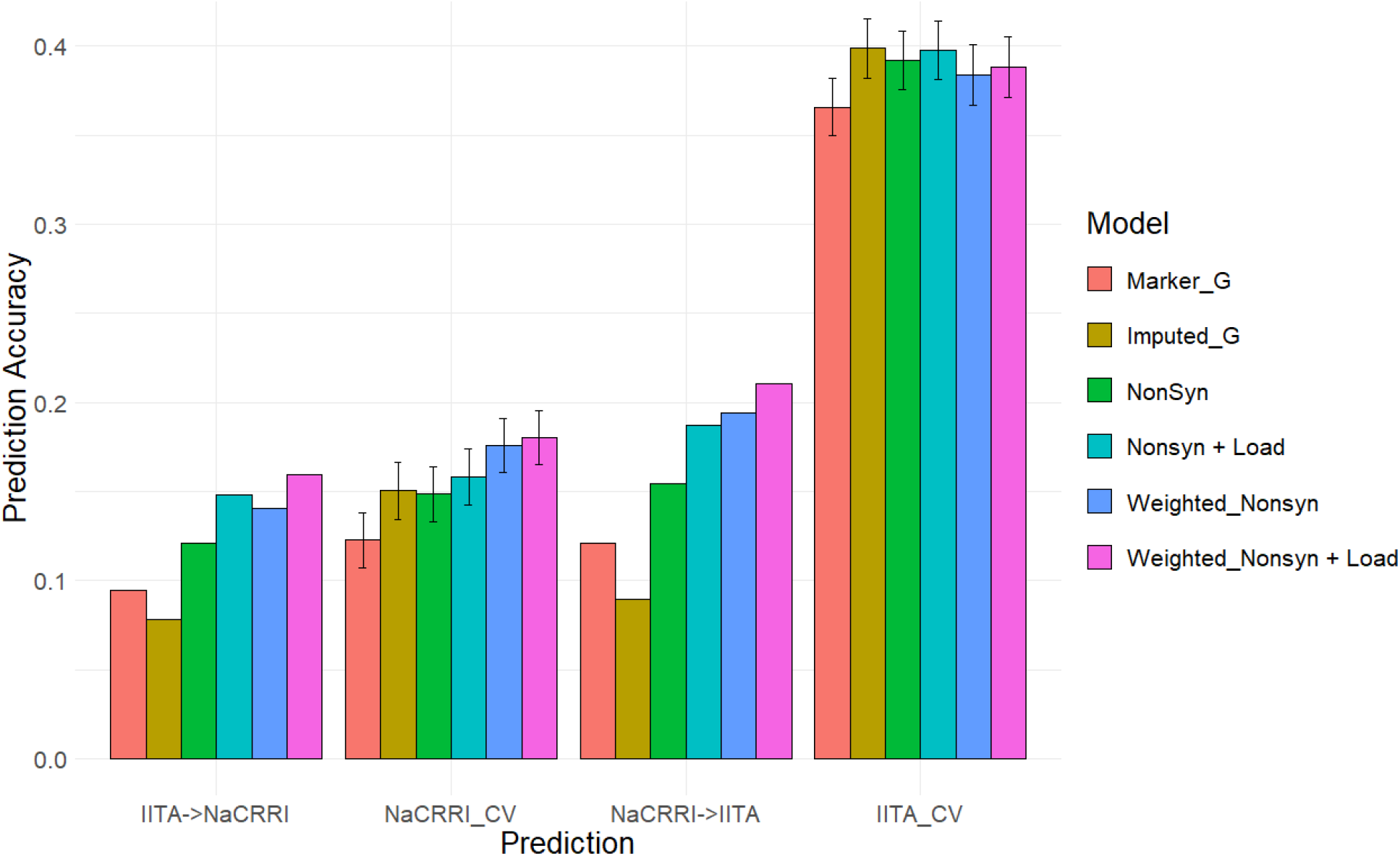
Fresh Root Yield Genomic Prediction Leveraging Deleterious Annotations. Prediction accuracy is measured in cross-population and within-population prediction scenarios. Genomic models are represented as bar graph colors where various genomic and deleterious data are used in the genomic prediction. Error bars represent a 95% confidence interval for within-population 10-fold prediction.

**Figure 7.**
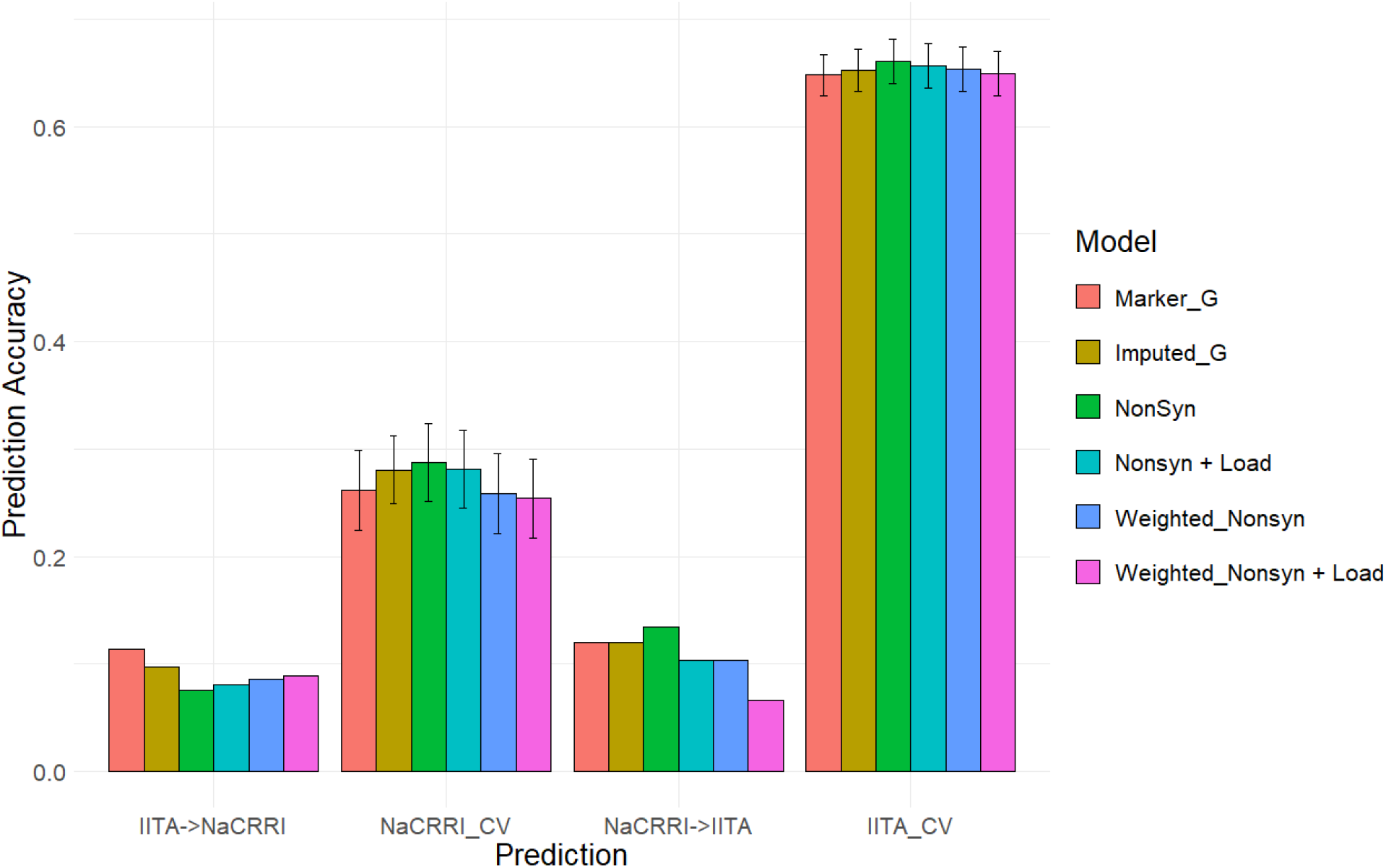
Dry Matter Percentage Genomic Prediction Leveraging Deleterious Annotations. Prediction accuracy is measured in cross-population and within-population prediction scenarios. Genomic models are represented as bar graph colors where various genomic and deleterious data are used in the genomic prediction. Error bars represent a 95% confidence interval for within-population 10-fold prediction.

## Discussion

Genetic load, as defined as the accumulation of deleterious mutations through domestication, drift, mutation-selection balance and other means, has been identified as an impediment to the genetic value of a crop (Agrawal and Whitlock, 2012; Smýkal et al., 2018). Through simulation, we explored the possible scenarios in which knowing the exact deleterious mutations could improve breeding selections. In this study, we went on to use evolutionary conservation to and genomic information to quantify deleterious mutations in cassava clones, as well as predict their potential effects.

### Simulation Informs Genomic Prediction Potential

The simulation of phenotypes under differing genetic architectures allowed us to manage expectations for the best possible scenarios in which understanding the causative variation of a trait could help inform genomic selection decisions. As we only observed benefits to genomic prediction under scenarios with <∼100 QTL, it is clear that LD structure captured by genome wide markers is sufficient for genomic prediction under highly complex genetic architectures (Fig 3). The scenario with the fewest QTL (<∼10) represents a more Mendelian or oligogenic architecture, which might benefit more from a marker assisted selection methodology, but it follows that traits with higher effect sizes of QTL will see more improvements from causative knowledge in genomic prediction. Interestingly, within-population predictions showed smaller, but still substantial benefits in genomic prediction accuracy. These results indicate that our empirical predictions have the potential to benefit from deleterious mutation annotations, only if there are a few or intermediate number of QTL (<∼100) with substantial effects. Importantly, the expected benefits shown through simulations depend directly upon the population and LD structure in our tested clones and cannot necessarily be useful to interpret potential benefits in other scenarios.

### Evolution Conservation Reveals Deleterious Mutations

We used evolutionary conservation and protein annotations to classify certain mutations as deleterious. By aligning over 50 species of relatively recent ancestry, we were able to assess the conservation status of a large majority of the cassava genome. We used separate neutral trees for each gene, rather than the entire chromosome or species, to address the difference between gene ancestry common in plants due to historical gene and genome duplication. Because of the millions of years of evolution, it is very difficult to predict the sizes of selection coefficients from evolutionary conservation alone (Huber et al., 2020). We then needed predicted protein effects of these mutations from SIFT to refine our set of putatively deleterious mutations. After defining our deleterious alleles, we separated the assessment of deleterious load into homozygous and heterozygous, because most deleterious mutations are assumed to be recessive (Bosse et al., 2019) and cassava has been shown to mask deleterious mutations through heterozygosity (Ramu et al., 2017). These assessments of genetic load are at least partially validated by a negative correlation between plant yield and homozygous deleterious mutations (Sup Fig 3.).

As previously mentioned, evolutionary conservation alone cannot easily resolve effect sizes of mutations. For this reason, we used protein perturbation information from SIFT and UniRep to prioritize functional variants similar to work recently done in Maize (Ramstein and Buckler, 2022). Another advantage of this weighting method is that it does not imply a directional effect of the mutations, thereby allowing for potential positive or adaptive effects (Loewe and Hill, 2010) of derived mutations at conserved sites.

### Leveraging Functional Data in Genomic Prediction

The inclusion of deleterious and functional mutations derived from evolutionary conservation showed promising value in informing the genetic value of cassava clones. Our results displayed improvements for cross-population predictions of fresh root yield as well as some of the within-population predictions in NaCRRI (Fig 6). This follows with the understanding that total plant growth, and even root yield, are correlated with total plant fitness (Pan and Price, 2001), while root dry matter percentage, which is primarily a quality trait, likely has little direct correlation with evolutionary fitness (Fig 7). We expect this trend would continue for other traits; however, few traits are measured identically across multiple populations.

In this study, we used multi-kernel GBLUP methods of genomic prediction to partition the additive and dominant genetic effects, while substituting unweighted and weighted genomic relationship matrices formed from subsets of the genomic data. These methodologies rely on the assumption that our selected functional variants, and the weights prescribed to them, are derived from a separate, and more functional, distribution of effects from a default, genome-wide relationship. Other methods, including Bayesian models, exist to prioritize functional information in genomic prediction, however multiple studies have found it to be difficult to prescribe consistent, significant differences in prediction accuracy results between them and GBLUP models, and the specific benefit of one method or the other are often situational (Moghaddar et al., 2019; Khansefid et al., 2020; Cheruiyot et al., 2022).

### Reflections on Load

In an effort to improve cassava’s role as a reliable food source around the world, our results show the importance and potential of addressing the impact of genetic load. We used evolution and protein annotations to determine these deleterious mutations responsible for genetic load. It is important to note that, while the methods used in this study detected impactful deleterious variation across the genome, they ignore the many deleterious mutations likely found in regulatory regions of the genome.

The improvements made in genomic prediction validate the effects of these deleterious mutations and offer one possible avenue for their potential application. As observed in the within-population prediction of IITA, where prediction accuracy is higher and unaffected by our annotations, the application of this understanding of genetic load may not be beneficial in every breeding scenario, however cross-population prediction is not the only instance where deleterious information may prove informative. Rapid cycle recurrent selection, where generations of selection occur without phenotyping, could be another situation in which tracking functional information across the genome could improve genomic selection decisions.

In addition to genomic prediction scenarios, the understanding of the deleterious mutations responsible for genetic load in cassava could suggest alternative methods for crop improvement. Many crops today utilize hybrid breeding, where multiple groups of inbred parents are bred for use in creating a superior hybrid. Selecting on inbred individuals exposes recessive, or partially recessive, deleterious mutations, allowing them to be effectively purged in fewer generations. While difficulties due to severe inbreeding depression in cassava have hindered this genre of breeding, efforts being made in crops like potato show it’s potential in a crop burdened by heavy genetic load (Bachem et al., 2019). Doubled haploidization has been a common tool in some inbred crops, while historically difficult to implement in some crops like cassava, however newer implementations such as those reported from ScreenSys (https://www.screensys.eu) offer a possible method of producing enough viable embryos for crops with heavy inbreeding depression like cassava. (Nasti and Voytas, 2021). With the understanding of the extent to which deleterious mutations account for missed potential in cassava performance, further consideration for how to effectively purge genetic load will be needed.

Historical evolution and population genetics continues to shed light on our understanding of genomic functions, as seen in our study in cassava. We showed the utility of using evolutionary derived deleterious mutations to improve genomic prediction across cassava populations. Additionally, the genetic load was identified from <∼100 homozygous deleterious mutations per clone (Fig 4). This number of mutations could be the target of further improvement through gene editing or other means. In the future, as genome sequencing accelerates, coupled with our understanding of protein functions, we may be able to make targeted decisions to purge genetic load from cassava and advance genetic gains.

## Acknowledgements

We would like to acknowledge the many germplasm sources that contributed tissue for sequencing (Sup. Table 1.) including: the Denver Botanic Garden, Germplasm Resources Information Network, the Missouri Botanic Garden, the Montgomery Botanic Garden, the National Botanic Garden, the National Tropical Botanic Garden, The New York Botanic Garden, and the US Botanic Garden. Their support was essential in sampling the vast number of species used in this study. We would also like to thank IITA and NaCRRI for contribution of data that we used to cassavabase. In particular, we thank Peter Kulakow, Ismail Rabbi, and Prasad Peteti who were project leads at IITA and Robert Kawuki, a project leader at NaCRRI, and Chiedozie Egesi, overall project manager of the NextGen Cassava Project.

## Methods

### Euphorbiaceae sequencing & Assembly

We gathered a total of 52 related species, 26 of which we sequenced and assembled, to evaluate evolutionary conservation across the cassava genome. In order to maximize the amount of evolutionary time sampled, while maintaining reliable alignments to cassava, we sampled 26 species across the Euphorbiaceae family, to which cassava belongs. We then sequenced these species using Illumina NovaSeq-6000. Genome sizes were estimated using k-mer spectra in order to estimate sequence input coverage for assembly. Additional short-read sequences were downloaded from SRA corresponding to 11 unspecified Euphorbiaceae taxa (Liu et al., 2019). We then used a short-read sequence assembler MEGAHIT (Li et al.), with modified parameters of “-m 0.2 -t 10 --no-mercy --min-count 3 --k-min 31 --k-step 20” to create contig assemblies. We additionally obtained long-read sequences using PacBio Sequel II for 7 species among our sampled Euphorbiaceae taxa. These sequences were assembled using Hifiasm (Cheng et al., 2021) utilizing default settings. An additional 15 genome assemblies from other related species were downloaded from SRA and added to our assembled genomes resulting in a total of 52 species in all (Sup. Table 1).

### Sequence Alignment and Evolutionary Conservation

We used gene alignments from Cassava V7.1 gene annotations to the 52 species to extract homologous gene sequences for multiple sequence alignment. Gene transcripts were aligned using minimap2, and the best aligned region with >= 90% alignment length matching was retained as homologous coding sequences for each species were then extracted and aligned using MAFFT (Katoh et al., 2002) multiple sequence alignment. With a multiple sequence alignment for each gene, we then generated gene trees using RAxML (Stamatakis, 2014), and calculated evolutionary rates using baseml from the PAML (Yang, 2007) suite of tools. We then identified ancestral alleles at every site across the genic regions of the genome, using the ancestral node containing Manihot, Hevea, and Cnidoscolus genera. We used evolutionary conservation to select representative gene models for each gene, as well as only retaining genes with 5’ and 3’ untranslated regions annotated resulting in ∼25k genes models (Sup. Table 2).

### Deleterious Mutations

We used evolutionary conservations & protein structure conservation to identify deleterious mutations and produce weights for functional importance of sites across the cassava Genome. Deleterious mutations were categorized as sites with a baseml evolutionary rate of <0.5 and a “Sorting Intolerant From Tolerant” (SIFT) score of < 0.05. Additionally, we required deleterious sites to have < 20% minor allele frequency in the cassava HapMap (Ramu et al., 2017) (Fig 4).

In addition to identifying a binary classification of deleterious, we used a RandomForest model to obtain a quantitative prediction of conservation similar to a previously reported method reported by Ramstein & Buckler (Ramstein and Buckler, 2022). We used baseml evolutionary rates to classify nonsynonymous sites as either conserved (evolutionary rate < 0.3) or non-conserved (evolutionary rate > 2), while sites with values outside these ranges were excluded from model training. SIFT, UniRep, and 100bp windowed GC% as predictors in the RandomForest model implemented by that R package “ranger” (Wright and Ziegler, 2017). From the SIFT database, we used both the mutation type and SIFT score, which gives the predicted deleterious effect a base-pair substitution. UniRep is a deep learning technique which characterizes protein structure (Alley et al., 2019), which we used to produce 256-unit representations of each protein and its associated mutated forms (https://github.com/churchlab/UniRep).

To increase the number of observations in the model, we used both the known HapMap mutations and *in silico* non-synonymous mutations at every possible site in our gene models. This resulted in over 1 million non-synonymous mutations whose genomic conservation could be modeled. We then used a leave-one-out prediction scheme where each of the 18 cassava chromosomes was left out of model training and predicted by the other 17. This method produced a predicted value between 0-1 for each of the ∼66k nonsynonymous, segregating mutations used in this study (Fig. 5).

### Phenotypic & Genotypic Data

Phenotypic and genotypic data for 1048 cassava clones were downloaded from *cassavabase*.*org* representing two populations of breeding lines. The first population is from a breeding program at International Institute of Tropical Agriculture (IITA) in Nigeria, while the second is from a breeding program at National Crops Resources Research Institute National Crops Resources Research Institute (NaCRRI) in Uganda, representing breeding material for West and East Africa, respectively. Genotypes for the associated clones were downloaded from the “*East Africa Clones Dart-GBS 2020*” genotyping protocol on cassavabase.org containing 23,431 variants. Plant phenotypes for fresh root yield and dry matter percentage were downloaded from *cassavabase*.*org* and prepared according to previously described methods (https://wolfemd.github.io/GenomicSelectionManual/index.html).

We then performed genotype imputation using the cassava haplotype map using Beagle5 (Browning et al., 2018), with an **Ne=100**, resulting in ∼26M variants. These variants were then filtered down to two genome-wide marker sets, one being a thinned sample of ∼135k genome-wide SNPs, and the other being all non-synonymous sites segregating in both populations resulting in ∼66k genome-wide variants. The input marker genotypes, the imputed sample, and the imputed non-synonymous sites will be used in genomic prediction analyses.

### Causative Variation Simulation

We used quantitative trait loci (QTL) simulation, replicated 50 times, to model the potential benefits of knowing causative variants in genomic prediction. This simulation begins by sampling QTL across the 66K variant sites from a binomial distribution with the probability of being a QTL varied across possible values of 10^−1^, 10^−2^, 10^−3^, and 10^−4^. The effect sizes for these QTL were then sampled from a gamma distribution using the *rgamma* function in R, with the shape parameter=1, with the ancestral allele set as having a positive effect. Lastly a dominance effect for each QTL was sampled from normal distribution rnorm(mean = 2,sd=0.3), restricting to dominance <=2 (Sup Fig 1). Phenotypes were then generated for the 1048 cassava clones. Residuals were then simulated such that the trait had a heritability of approximately 0.3.

### Genomic Prediction Models in Simulation

We performed cross-population and 10-Fold within-population predictions using the simulated data, with and without QTL information incorporated into the prediction model. Genomic prediction was performed by using GBLUP methods fit using ASReml, with additive and dominance effects modeled as separate kernels. For all models described, residuals are represented by ε and modeled as random with **ε**∼N(**0**,**I**σ^ε2^).

For prediction using simulated phenotypes, we compared three different models.

The first model represents our baseline prediction:

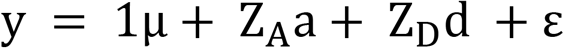

Where ŷ is the simulated phenotype entries, mu is the phenotype mean, **a** is the vector of additive genetic effects, **Z**_**a**_ is the incidence matrix, and **a**∼N(**0**,**G**_**A**_σ_a_^2^), G_A_ is an additive genomic relationship matrix produced using the VanRaden (VanRaden, 2008) method, and σ_a_^2^ is the additive genetic variance.

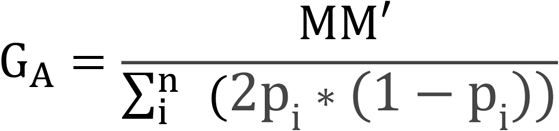

Where M is the scaled genotype matrix (where genotypes are stored as dosages of 0,1,and 2 referring to being homozygous for reference allele, heterozygous, and homozygous for the alternate allele, respectively) and p_i_ is and allele frequency at the ith locus. **Z**_**D**_ and **d** are analogous to the additive method, with the exception that a dominance genomic relationship matrix is produced using the Nishio and Satoh (Nishio and Satoh, 2014) method.

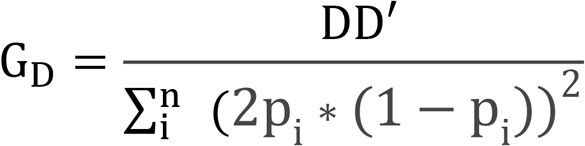

Where the entries of D are given as -2p_i_^2^ for the homozygous reference allele, 2p_i_*(1-p_i_) for the heterozygote, and 2(1-p_i_)^2^ for the homozygous alternate allele.

The second model includes additive and dominance QTL relationship matrices formed in identical manner to the **G**_**A &**_ **G**_**d**_ matrices, but only utilizing the known QTL sites in the genomic relationship matrices:

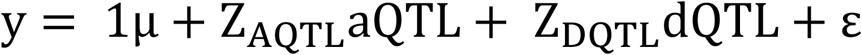

The final model includes weighted QTL matrices based on their effect size:

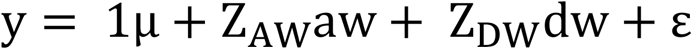

Here the weighted matrices are formed using modified methods of the previously cited methods. The weighted additive matrix given by:

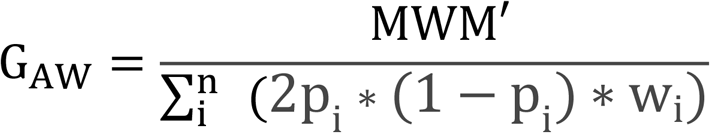

Where M is the scaled genotype matrix. W is a diagonal matrix with w_i_ along the diagonal, w_i_ and p_i_ are the weight and frequency for the ith locus, respectively.

The Weighted Dominance matrix is modified in a similar fashion to the additive matrix:

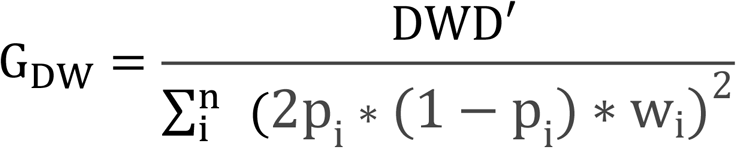

Where the entries of D are given as -2p_i_^2^ for the homozygous reference allele, 2p_i_*(1-p_i_) for the heterozygote, and 2(1-p_i_)^2^ for the homozygous alternate allele.

### Genomic Prediction Models in Empirical Data

The genomic prediction models used for real breeding program phenotypes follow a similar pattern to our simulated scenario, with a few notable differences.

First, our ground truth for the phenotype of each clone was the best linear unbiased estimate (BLUE) using a model similar to those previously used in cassava plot level traits (Wolfe et al., 2017) and those suggested for use with African cassava breeding data (https://wolfemd.github.io/GenomicSelectionManual/index.html):

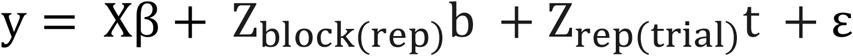

where y is the vector of the phenotype, β included a vector of fixed effects for the population mean, the location–year combination, the number of plants harvested per plot, and germplasm ID with design matrix **X**. Replications were nested in trials, treated as random, and represented by the design matrix **Z**_**rep(trial)**_ and the effects vector **t**∼N(**0**,**I**σ_t_ ^2^). Blocks were nested in replications, treated as random, and represented by the design matrix **Z**_**block(rep)**_ and the effects vector **b**∼N(**0**,**I**σ_b_ ^2^).

Having a ground truth phenotype, we then compared multiple different genomic prediction models to measure the potential benefits to including the deleterious annotations. Each model followed a similar form:

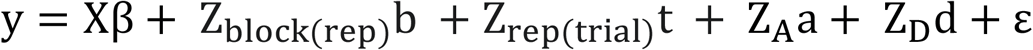

This generic model mirrors the previous one, with the exception that germplasm ID is no longer treated as fixed but is instead **Z**_**A**_ and **Z**_**D**_ are design matrices indicating observations of germplasm IDs for the vectors of additive and dominance effects **a** and **d**, modeled as previously described in the simulated scenario. The six models we compared involve substituting different markers and methods of constructing genomic relationship matrices for **Z**_**A**_ and **Z**_**D**_, as well as adding fixed effects for derived homozygous and heterozygous load. The six models include:

- *Marker_G* where the 23,431 variants are used to produce the genomic relationship matrices.
- *Imputed_G* where ∼135k imputed genome-wide segregating sites are used to produce the genomic relationship matrices.
- *Nonsyn* where 66k imputed, segregating, nonsynonymous mutation sites are used to produce the genomic relationship matrices.
- *Nonsyn + Load* which is identical to *Nonsyn* with the exception of including the derived load as fixed effects in the prediction
- *Weighted_Nonsyn* uses the same sites as *Nonsyn*, however the genomic relationship matrices are created using the weighted method described previously, with the deleterious weights for each SNP.
- *Weighted_Nonsyn + Load* which is identical to the *Weighted_Nonsyn* with the exception of including the derived load as fixed effects in the prediction

Each model was evaluated by performing the cross-population and within-population predictions as previously described and using the correlation between predicted phenotype and the BLUE as the prediction accuracy (Fig 6&7). Prediction accuracy was also calculated as the number of the top 25 performing clones predicted as being among the top 25 performing clones (Sup Fig 6&7.).

## Data Availability

Genotype and Phenotype data used in this study is available at cassavabase.org. Euphorbiaceae sequence reads and assemblies generated in this study will be available under PRJEB55682 on the European Nucleotide Archive. Code used to process data and produce assemblies, simulations, genomic predictions are available at https://bitbucket.org/bucklerlab/cassava_load_and_gp

## Supplemental Data

### Supplemental Figures

**Supplementary Figure 1.**
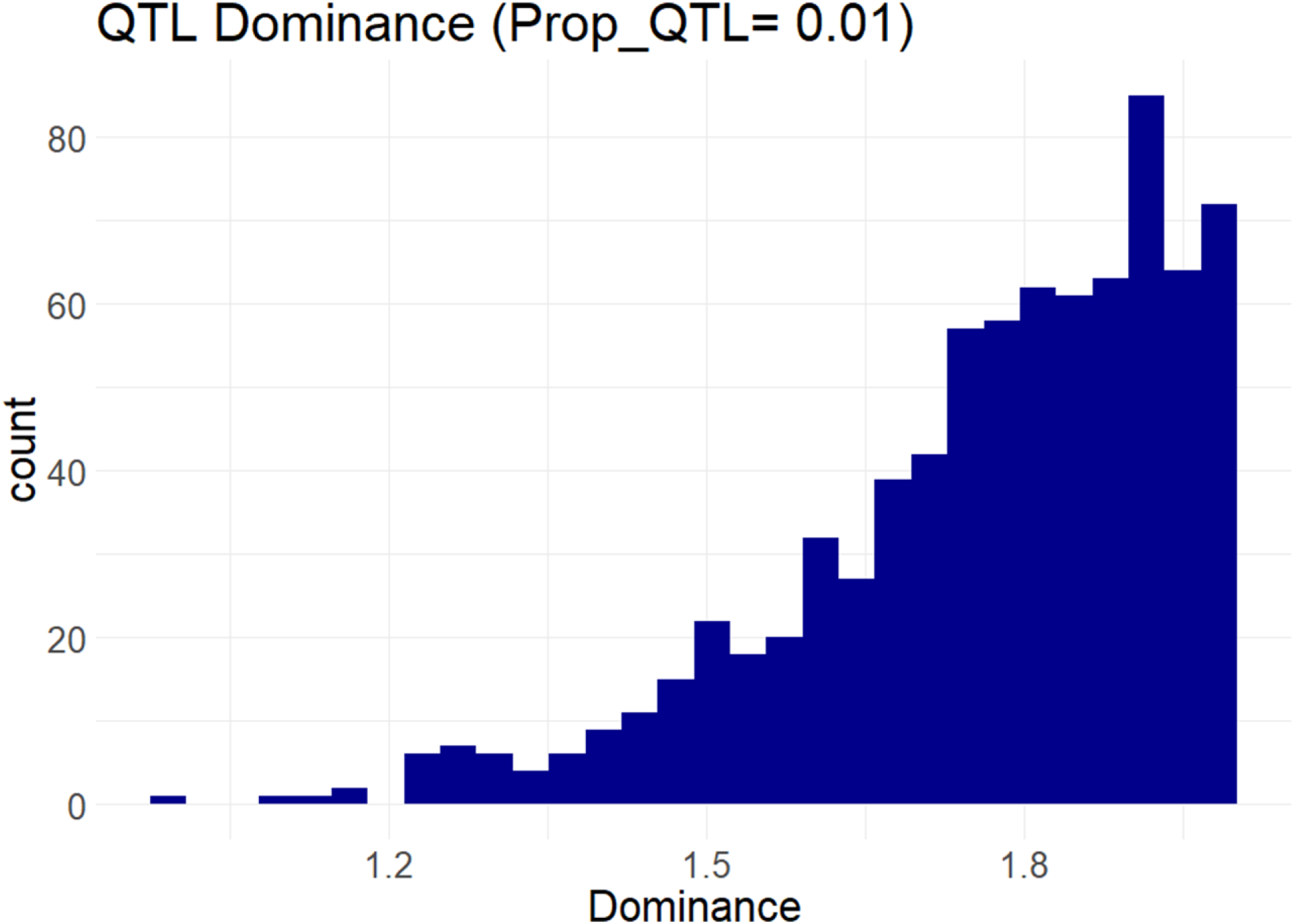
QTL Dominance. Distribution of QTL dominance from one simulation instance. A dominance value of 2 indicates a QTL as completely dominant where the value of heterozygous genotype is equal to a homozygous genotype, while a value of 1 indicates a QTL is completely additive, where the value of a heterozygous genotype is half the value of a homozygous genotype.

**Supplementary Figure 2.**
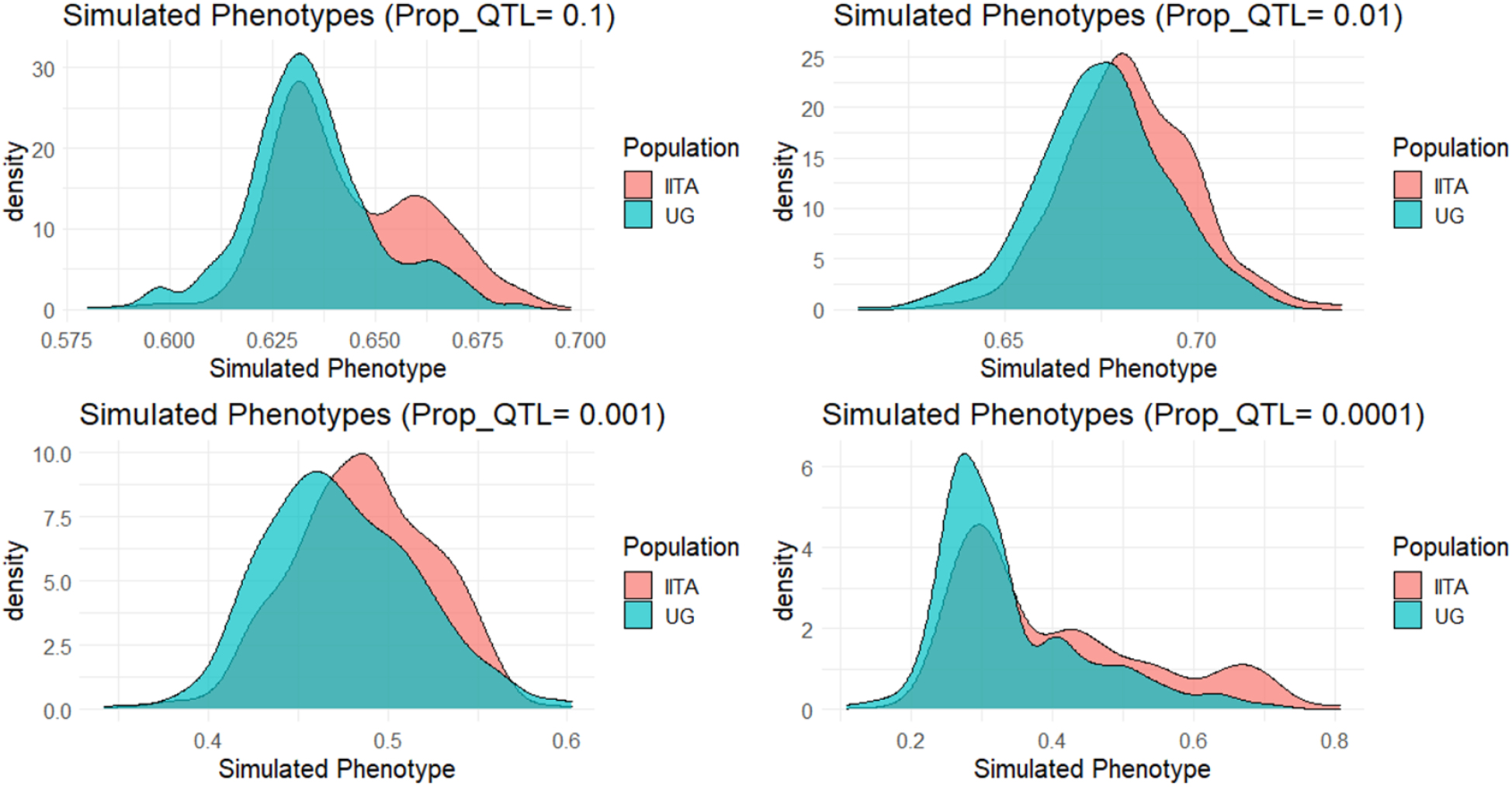
Simulated Phenotypes. Phenotypes simulated under one example simulation instance for each genetic architecture.

**Supplementary Figure 3.**
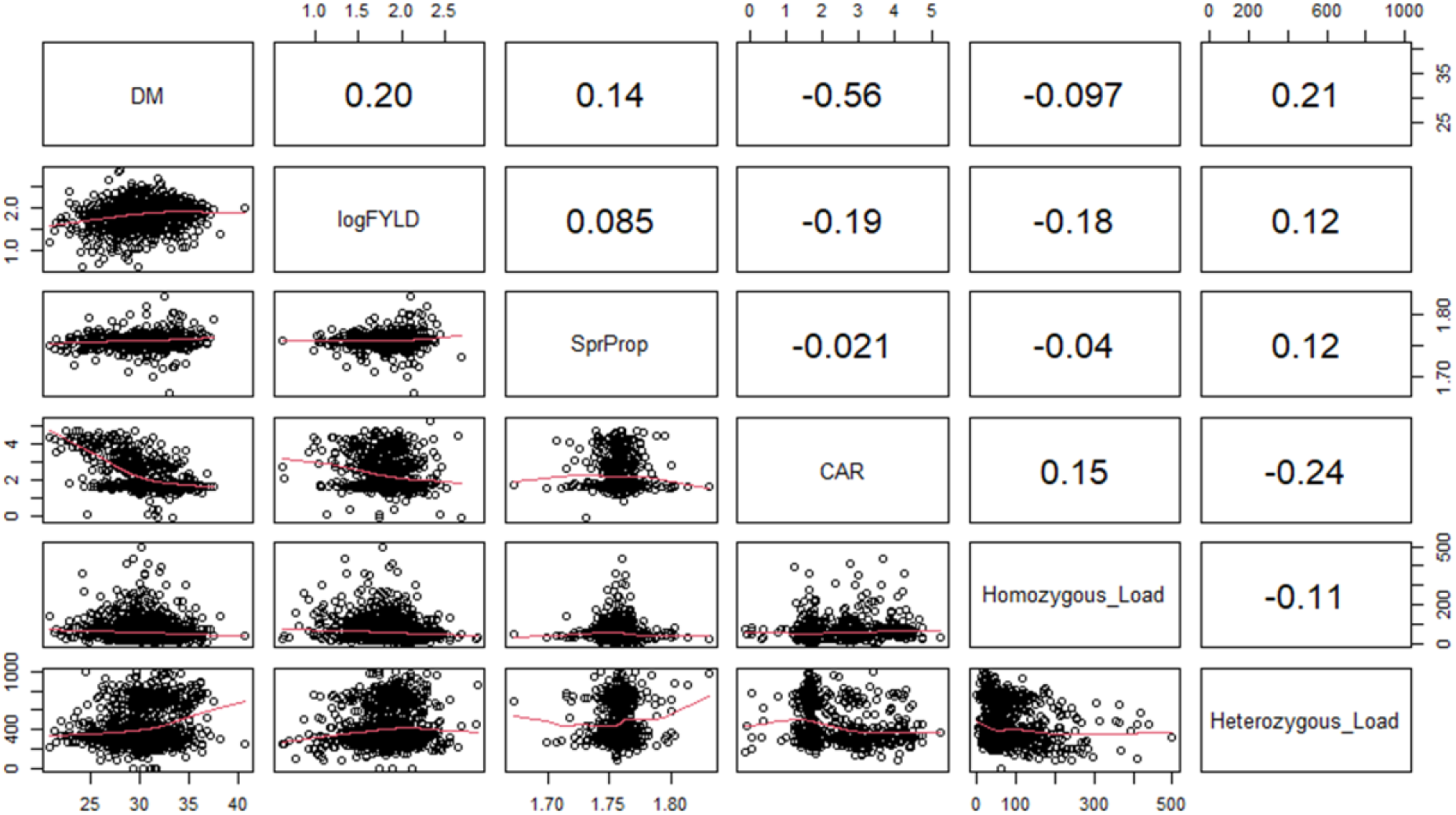
Trait and Deleterious Mutation Correlations. Correlations between traits: dry matter percentage (DM), log transformed fresh root yield (logFYLD), proportion of viable sprouts (SprProp), and carotenoid content (CAR). Correlation to homozygous and heterozygous genetic load measured by number of derived deleterious alleles.

**Supplementary Figure 4.**
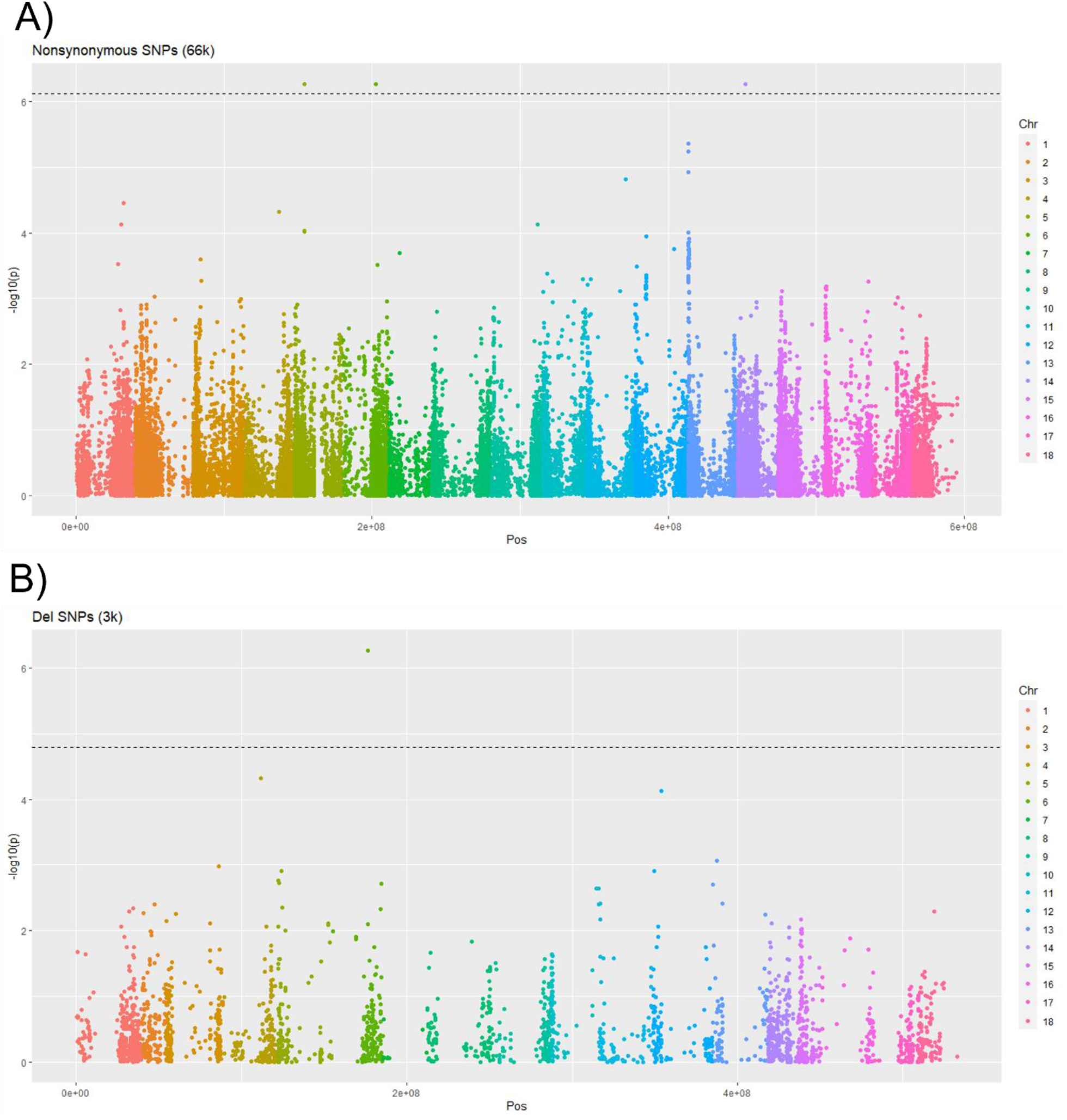
Genome Wide Association Fresh Root Yield. Genome wide association using Tassel5 MLM model y∼G+PCs(1-5). A) Association performed on 66k Nonsynonymous SNPs, where 5 variants at 3 loci passed Bonferroni significance. B) Association performed on ∼3k deleterious SNPs, where 1 variant passing Bonferroni significance.

**Supplementary Figure 5.**
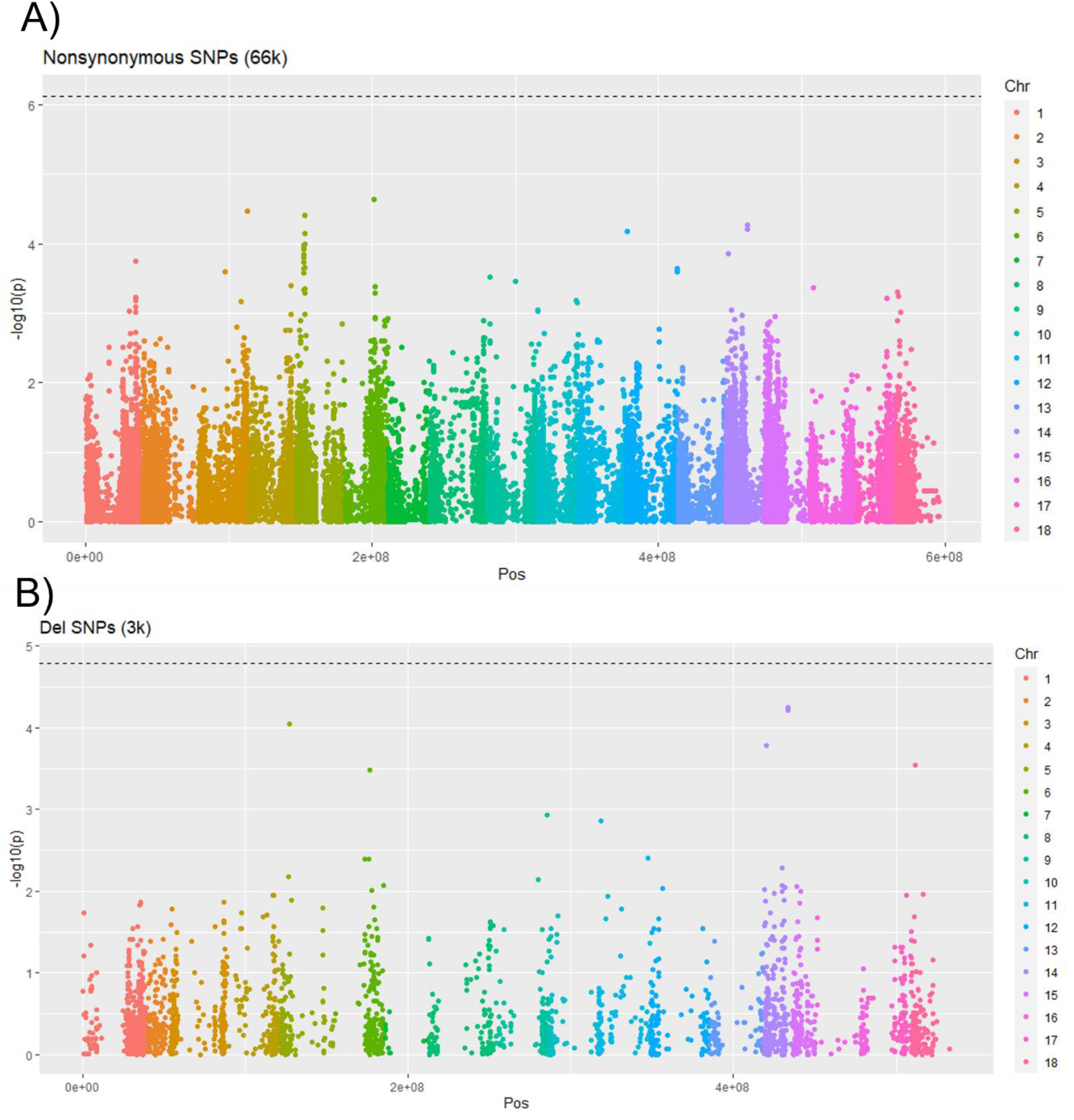
Genome Wide Association Dry Matter Percentage. Genome wide association using Tassel5 MLM model y∼G+PCs(1-5). A) Association performed on 66k Nonsynonymous SNPs. B) Association performed on ∼3k deleterious SNPs.

**Supplementary Figure 6.**
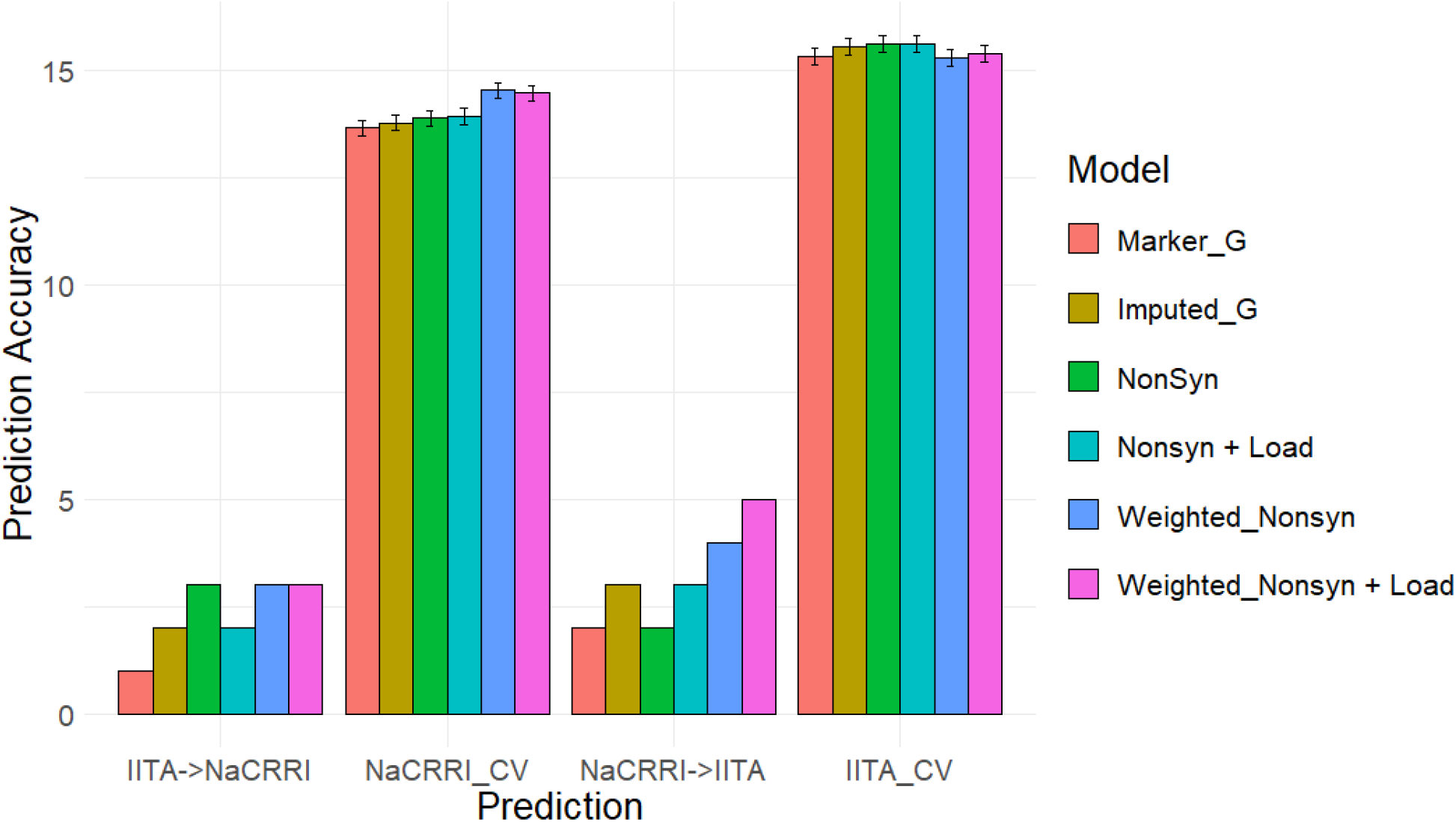
Top 25 Prediction Accuracy for Fresh Root Yield. Prediction accuracy is measured by number of 25 predicted clones that are among the top 25 performing clones. Genomic models are represented as bar graph colors where various genomic and deleterious data are used in the genomic prediction. Error bars represent a 95% confidence interval for within-population 10-fold prediction.

**Supplementary Figure 7.**
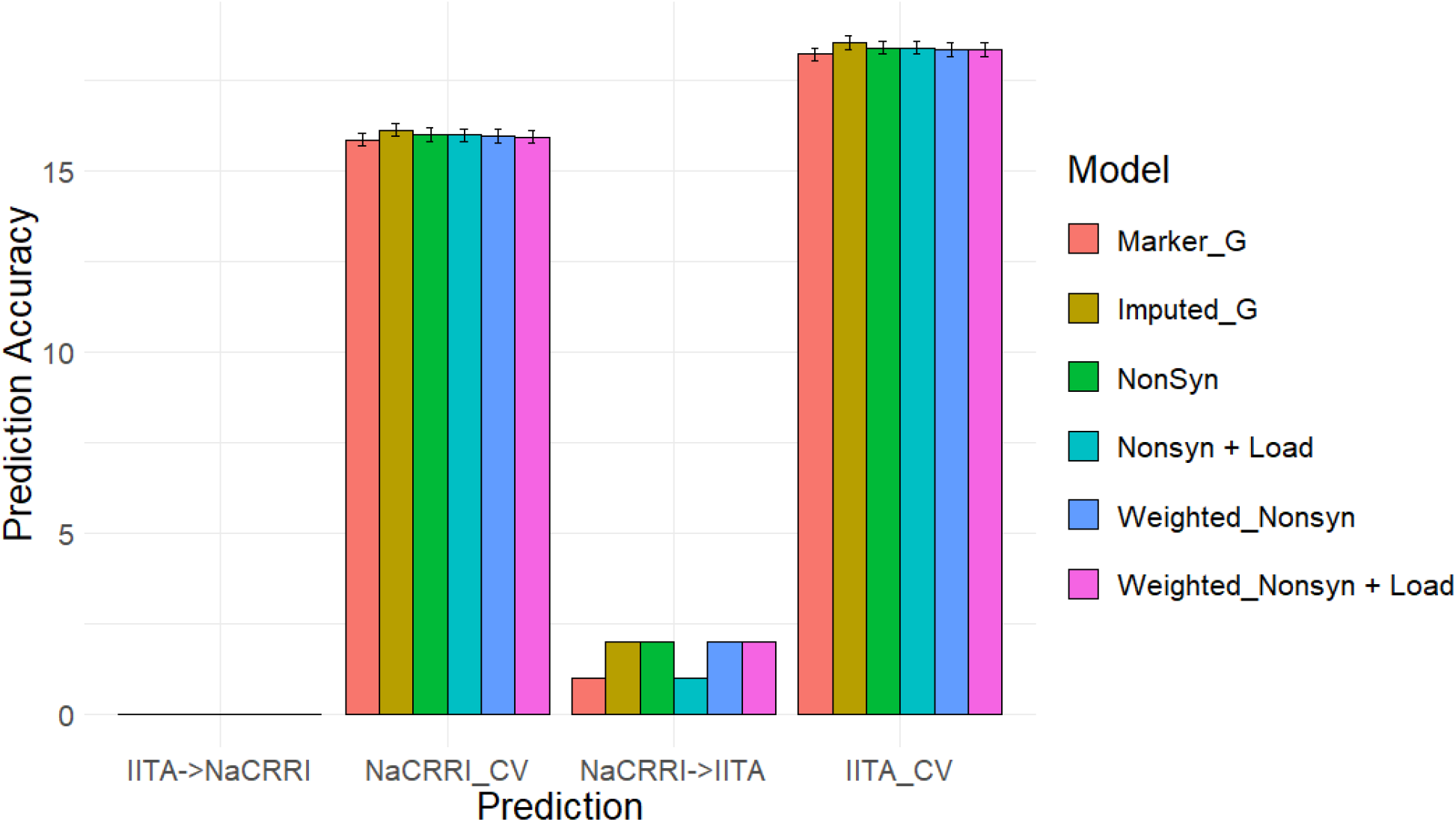
Top 25 Prediction Accuracy for Dry Matter percentage. Prediction accuracy is measured by number of 25 predicted clones that are among the top 25 performing clones. Genomic models are represented as bar graph colors where various genomic and deleterious data are used in the genomic prediction. Error bars represent a 95% confidence interval for within-population 10-fold prediction.

